# A parasitic flagellate among the Nucleariae: a new perspective on the phylogeny of Holomycota (Opisthokonta)

**DOI:** 10.64898/2026.07.21.739728

**Authors:** Alexei O. Seliuk, Igor R. Pozdnyakov, Andrey E. Vishnyakov, Darya O. Drachko, Sergey A. Karpov

## Abstract

Holomycota, together with Holozoa, represents one of the two main branches of the supergroup Opisthokonta. Fungi, as well as three basal protist lineages Nuclearida, Rosellida, and Aphelida belong to Holomycota. Surprisingly, among all these holomycotan groups, only the most basal lineage, Nuclearida, lacks any traces of flagella in its life cycle and is represented by free-living amoebae, in contrast to the obligately parasitic and flagellated Rosellida and Aphelida, which evolved later. As the earliest-branching lineage, nucleariids play a key role in understanding the origin and diversification of Opisthokonta, which is presumed to have evolved from a uniflagellated ancestor. In this study, we describe the first known nucleariid that is parasitic on algae and possesses a flagellated stage in its life cycle. We studied the life cycle, and showed the ultrastructure of main life cycle stages of this parasite. The multigene phylogenetic tree based on the newly sequenced transcriptome demonstrated its basal position among nucleariids, forming a clade of environmental OTUs. This finding together with unique ultrastructural features, parasitic style of life, and presence of flagellum allowed us to describe a new species, genus, and family in the order Nuclearidida. We emended the diagnoses of the order, class, and phylum Nuclearida. Based on the ribosomal phylogenetic tree where the *Insolitus nuclearius* has a basal position among the nucleariids we discussed the evolution of this group from the uniflagellated ancestor and provided new insights into the early evolution of Holomycota in a whole, revising hypotheses regarding the opisthokont ancestor.

## Introduction

Endo- and ectoparasites of algae are widely represented across the eukaryotic phylogenetic tree. They are distributed among different supergroups, including Alveolata, Rhizaria, Amoebozoa, and Opisthokonta. These parasites play a crucial role in aquatic ecosystems by recycling the cellular contents of large algae, which are otherwise inedible for planktonic invertebrates, back into the organic matter cycle (Kagami et al., 2014). Their role in regulating algal blooms is often underestimated: algal parasites compete for hosts and can almost completely eliminate dominant algal species, thereby altering ecosystem dynamics (Reñé et al., 2022). Their ability to cause outbreaks in dense algal populations is detrimental to aquaculture in open systems and to laboratory collections (Gromov, 2000; Letcher et al., 2015).

Many biological characteristics of these parasites remain poorly understood. However, advances in molecular biology, including genomic and transcriptomic approaches, have significantly improved our understanding of the diversity, phylogenetic relationships, and life cycles of alveolate and opisthokont parasites. According to recent phylogenetic reconstructions, Nuclearida, Rosellomycota, and Aphelidiomycota diverged sequentially from the ancestral lineage leading to the “true” fungi (Strassert and Monaghan, 2022; Mikhailov et al., 2022; Aleoshin et al., 2026). True fungi lack phagotrophic nutrition, and the transition to exclusively osmotrophic nutrition coincided with the early loss of genes associated with phagotrophy (Mikhailov et al., 2022; Pozdnyakov et al., 2023, 2025; Moreira et al., 2025). Despite differences in nutrition, many mycologists now include rosellids and aphelids within the kingdom Fungi, based on similarities in their life cycles to those of zoosporic fungi (Wijayawardene et al., 2024, 2025).

Aphelids and rosellids are intracellular parasites that phagocytize host cell contents, whereas nucleariids are heterotrophic, free-living organisms. This relatively small group of amoebae consists of spherical cells bearing radiating filopodia and sometimes containing two or more nuclei (Gabaldon et al., 2022). Nucleariids represent the earliest-branching lineage within Holomycota and thus play a key role in understanding the origin and diversification of Opisthokonta.

In this study, we report the first known nucleariid that is parasitic on algae and possesses a flagellated stage in its life cycle. This finding helps to explain how nucleariids—previously considered exclusively amoeboid—form the lineage closest to the opisthokont ancestor, which is inferred to have been uniflagellated. It also provides a new perspective on the origin and evolution of both Holomycota and Holozoa.

## Materials and Methods

### Isolation and cultivation

Strain X-141 was isolated from the marine sample Cl-4, which also served as a source of *Protaphelidium rhizoclonii* (Seliuk and Karpov, 2024). Cultivation was carried out following the protocol described in Seliuk and Karpov (2024), with the exception that *Rhizoclonium* sp. was grown in Petri dishes containing a thin layer of 1% agar prepared with 25‰ artificial seawater.

### Light microscopy

Microscopic observations of living cultures were performed using Leica DM2500 or Nikon Eclipse Ni upright microscopes equipped with DIC and phase-contrast optics and a DS-Fi-3 camera (Nikon, USA) controlled by NIS-Elements AR software (Nikon). For each observation, several infected algal filaments were transferred into a small droplet of medium taken from a Petri dish. In some cases, such droplets were maintained for several hours or days in a humid chamber.

For prolonged observations, time-lapse imaging was performed using an inverted Leica DM3000 microscope equipped with DIC and phase-contrast optics and LAS X software (Leica Microsystems, Germany). For this purpose, special Petri dishes with a thin bottom were used.

### Transmission (TEM) and scanning (SEM) electron microscopy

For TEM, infected algal filaments were fixed in 2% glutaraldehyde in 0.1 M cacodylate buffer for 4 h at 4 °C. The fixative containing zoospores was then filtered through a 13-mm polycarbonate filter with a 1-μm pore size (ipPore™ Track-Etched Membrane, Belgium) using a Swinny Filter Holder (13 mm; EMD MilliporeSigma, USA). The remaining algal material was collected using a Pasteur pipette, transferred onto the filter in a small droplet, and pressed down by filtration. The fixed material on the filter was embedded in 2% low-melting-point agarose. Agarose blocks were washed in 0.1 M cacodylate buffer for 15 min and post-fixed in 1% OsO₄ in the same buffer for 1 h at 4 °C.

Subsequently, the samples were washed again, dehydrated through a graded ethanol series followed by propylene oxide, and embedded in Agar LV resin (Agar Scientific Ltd., UK). Ultrathin sections were prepared using a Leica Ultracut ultramicrotome (Leica, Germany) with a diamond knife. After double staining with uranyl acetate and lead citrate, sections were examined using a JEM 1400 transmission electron microscope (JEOL, Japan) equipped with an Olympus Veleta digital camera (Olympus, Japan).

For SEM, infected algal filaments and culture containing swimming zoospores were transferred onto coverslips pre-cleaned with 70% ethanol and left for 10 min to allow zoospores to attach. An equal volume of 4% glutaraldehyde in 0.1 M cacodylate buffer was then added to achieve a final concentration of 2%, and the samples were incubated in a humid chamber for 30 min. After fixation, samples were washed with buffer, dehydrated through an ethanol series, and air-dried in a laminar flow cabinet. Specimens were mounted on aluminum stubs, coated with gold–palladium, and observed using a Hitachi TM-1000 microscope at an accelerating voltage of 15 kV.

### DNA amplification and sequencing

Individual zoospores were transferred into 0.2 μl PCR tubes containing 1 μl of medium using a micromanipulator. For PCR amplification, 10–50 zoospores were collected per tube, whereas 200 zoospores per tube were used for next-generation sequencing (NGS), after which samples were frozen at −80 °C.

To obtain ribosomal operon sequences, DNA was extracted using the PicoPure kit (Thermo Fisher Scientific, USA) following the manufacturer’s protocol, and 1 μl of the extract was used as a template for PCR. The first half of the 18S rDNA gene was amplified using primers RibA and S14R, and the second half using primers S.12.2 and RibB. The ITS region was amplified using OpistOut and 28r1 primers. All PCR reactions were performed using the Tersus Plus PCR kit with Tersus polymerase (Evrogen, Russia).

The PCR program consisted of an initial denaturation at 94 °C for 5 min; followed by 35 cycles of denaturation at 94 °C for 15 s, annealing at 60 °C for 30 s, and extension at 72 °C for 1 min; with a final extension at 72 °C for 5 min. Sequencing was performed on an ABI Prism 3500xl sequencer (Applied Biosystems, USA) using the same primers. Chromatograms were analyzed and assembled into full-length sequences using UGENE (Okonechnikov et al., 2012).

### cDNA production and transcriptome assembly for multigene phylogenetic analysis

Frozen flagellated cells were used as material for transcriptome assembly of the new organism. cDNA was synthesized from mRNA using the NEBNext® Single Cell/Low Input RNA Library Prep Kit for Illumina (NEB #E6420) following the low-input protocol. The concentration of the resulting cDNA was measured using a Qubit fluorometer. The cDNA library was sequenced on an Illumina platform, yielding 76,877,119 paired-end reads of 150 bp.

The full-length ribosomal operon was extracted from the NGS data using RiboDetector v0.3.2 (Deng et al., 2022). The 18S rRNA gene sequence obtained from NGS data was compared with that obtained by Sanger sequencing and found to be identical. Therefore, the complete ribosomal operon derived from NGS data was deposited in GenBank instead of only the 18S rDNA sequence obtained by Sanger sequencing (accession number: XXX).

Read quality was assessed using FastQC, and low-quality regions were trimmed using Trimmomatic v0.40 (Bolger et al., 2014). Transcriptome assembly was performed using Trinity v2.15.2 (Grabherr et al., 2011). Assembly quality was evaluated using QUAST v5.0.2 (Gurevich et al., 2013), and completeness was assessed using BUSCO v5.4.2 with the single-copy ortholog database (Manni et al., 2021; Simão et al., 2015). The resulting transcripts were further filtered and validated using BlobToolKit 2 (Challis et al., 2020). BUSCO analysis indicated that the assembly was not sufficiently complete for publication of the full transcriptome; therefore, it was used only for multigene phylogenetic analysis.

### Ribosomal phylogenetic analysis

For phylogenetic analysis based on 18S rDNA, all available nucleariid sequences from GenBank were used, including the recently published sequence of *Nuclearia leuckarti* (Seliuk et al., 2024a), as well as additional environmental sequences from Aleoshin et al. (2026). Sequences were aligned using MAFFT v7.526 with the FFT-NS-2 algorithm (Katoh et al., 2013). Ambiguously aligned regions were removed using TrimAl v1.4.rev15 with the automated1 option (Capella-Gutiérrez et al., 2009). The final alignment consisted of 1,363 positions. GenBank accession numbers of all sequences are provided in Table S1.

The maximum likelihood (ML) tree was inferred using IQ-TREE 3 with 1,000 bootstrap replicates and the TIM+F+R2 substitution model selected automatically by ModelFinder (Wong et al., 2025). Bayesian inference was performed using MrBayes v3.2.7a under standard settings (Ronquist et al., 2012). The resulting trees were visualized using iTOL (Letunic and Bork, 2021).

### Multigene phylogenetic analysis

To construct the multigene phylogenetic tree, the transcriptome assembly of the studied organism was combined with genomic and proteomic datasets from 20 representatives of major Opisthokonta subgroups (NCBI accession numbers in parentheses).

Holomycota: *Fonticula alba* (GCA000388065.2), *Parvularia atlantis* (GCA943704415.1) (Nucleariida); *Rozella allomycis* (GCA000442015.1), *Encephalitozoon cuniculi* (GCA000091225.2), *Metchnikovella incurvata* (GCA003600395.1) (Rozellida); *Amoeboaphelidium protococcorum* (GCA023515745.1), *Aphelidium insulamus* (GCA036784855.1) (Aphelida); *Allomyces macrogynus* (GCA000151295.1), *Blastocladiella emersonii* (GCA025594325.1) (Blastocladiomycota); *Synchytrium endobioticum* (GCA006535955.1), *Spizellomyces punctatus* (GCF000182565.1).

Holozoa: *Sphaeroforma arctica* (GCA001186125.1), *Amoebidium appalachense* (GCA963693365.1), *Corallochytrium limacisporum* (GCA002811645.1) (Teretosporea); *Pigoraptor vietnamica* (GCA943704525.1), *Ministeria vibrans* (GCA943704425.1), *Capsaspora owczarzaki* (GCA000151315.2) (Filasterea); *Monosiga brevicollis* (GCA000002865.1), *Salpingoeca rosetta* (GCA000188695.1) (Choanozoa); *Sycon ciliatum* (GCA964019385.1) (Metazoa).

Additionally, three Amoebozoa species were included as an outgroup: *Dictyostelium discoideum* (GCA000004695.1), *Acanthamoeba castellanii* (GCA000313135.1), and *Vannella* sp. (GCA023345925.1).

Protein sequences corresponding to BUSCO ortholog sets for Metazoa and Fungi were extracted or translated using BUSCO v5.4.2. After removing duplicates, a dataset of 918 orthologous protein groups was obtained. Multiple sequence alignments were generated for each group using MAFFT v7.526, followed by trimming with TrimAl v1.4.rev15.

A preliminary IQ-TREE analysis was performed to assess sequence quality, and sequences failing the χ² test were removed. The remaining sequences were realigned and trimmed. Only alignments containing at least 75% of taxa (18 out of 24 species) were retained.

The final dataset consisted of 269 alignments corresponding to 269 single-copy orthologous genes. Phylogenetic reconstruction was performed using IQ-TREE 3 in concatenation and partitioned modes, with automatic model selection and 100,000 ultrafast bootstrap replicates.

## Results

### Light microscopy and SEM observations

The life cycle of the parasite includes the following stages. The dispersal stage is represented by a zoospore bearing a single posterior flagellum (Fig. 1A, Q). The zoospore settles on the algal surface and transforms into a cyst (Fig. 1J, K). Its contents penetrate the host cell wall with a pseudopodium and enter the algal cell (Fig. 1K, L). The intracellular amoeboid stage phagocytizes the host cell contents and subsequently divides into two daughter cells (Fig. 1M). These cells develop a flagellum and are released through a pre-existing or newly formed opening in the algal wall, or migrate into adjacent cells through transverse septa (Fig. 1N, O).

**Fig. 1.**
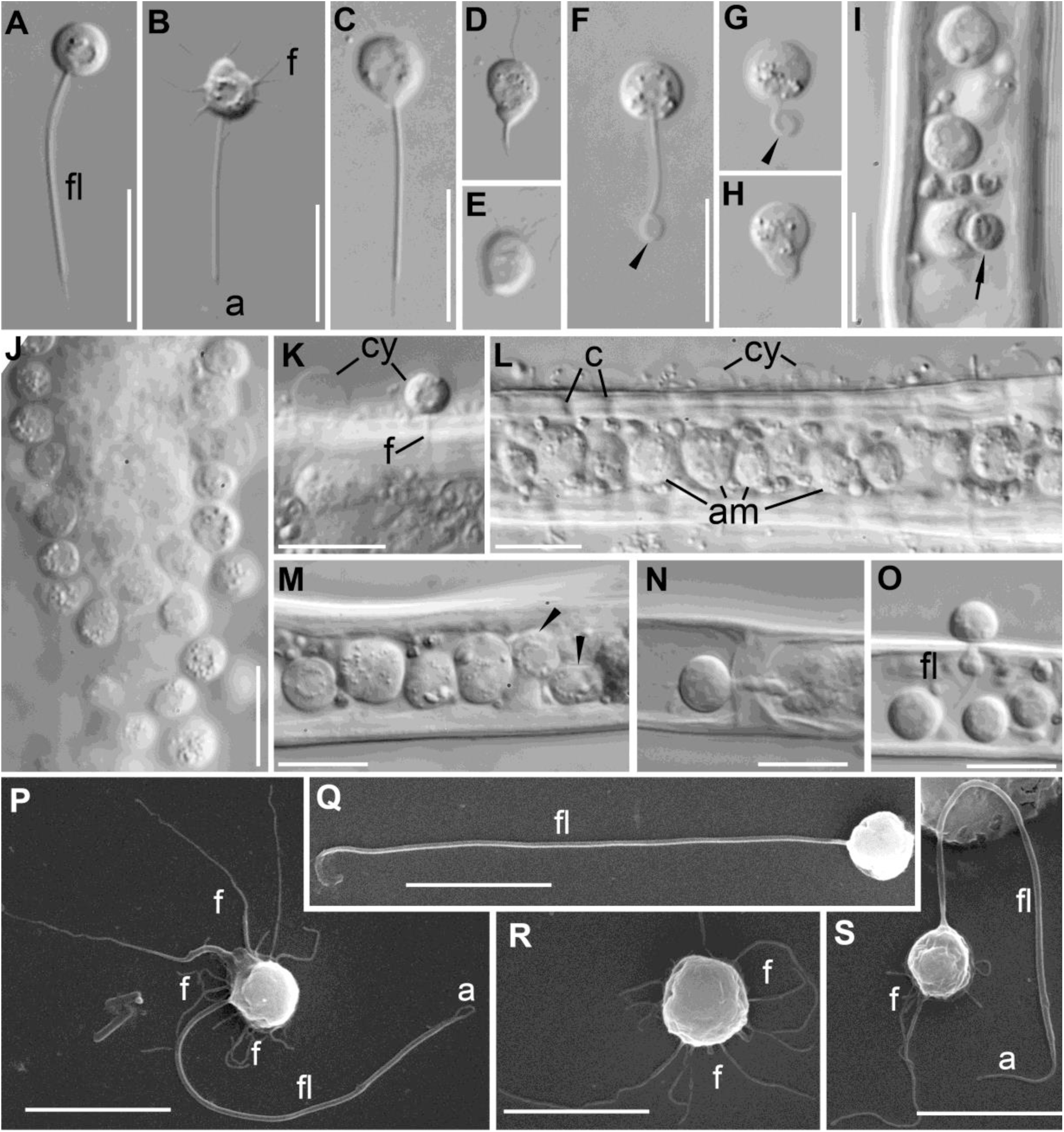
Light microscopy of the main life cycle stages of *Insolitus nuclearius*. A-O – LM images, P-S – SEM images. A-H – zoospores: A – flagellate, B – amoeboflagellate, C-E and F-H – two different types of flagellar retraction (arrowheads point a curl), I -excretion of big granule (arrow) from parasite, J – multiple infection of the host, K – encysted parasite penetrates the host cell wall, empty cyst to the left, L – eaten algal cell filled with vegetative cells of parasite, note perforated cell wall and empty cysts on its surface, M – two daughter cells (arrowheads) just after cell division, N – amoeboid cell moving to the left penetrates septa between algal cells, O – zoospore with posterior flagellum releases. P,S – amoeboflagellates, Q – flagellate, R – amoeba. Scale bars: A-J – 10 µm, K-S – 5 µm. **Abbreviations**: a – acroneme, am – amoebae, ax – axoneme, aw – algal wall, bf – basal fibers of non-flagellar kinetosome, bp – basal plate of kinetosome, с – сanal, cw – cyst wall, cy – cyst, er – endoplasmic reticulum, fk – flagellar kinetosome, fl – flagellum, fp – fibrillar plate at transition fiber attachment, fr – fibrillar root, fs – fibrillar sheath of non-flagellar kinetosome, ga – Golgi apparatus, l – lipid globules, m – mitochondrion, mb – microbody, mr-microtubular root, mt – microtubule, n – nucleus, nf – non-flagellar kinetosome, nu – nucleolus, ps – pseudopodium, st – starch, tp – terminal plate.

The flagellated zoospore is capable of forming branching filopodia, which it uses for movement while the flagellum remains non-motile (Fig. 1B, P, S). The cell body measures 4–5.5 µm in diameter, whereas the flagellum reaches up to 22 µm in length (including a 2.5 µm acroneme), corresponding to approximately 4–5 times the cell length. The flagellum is retracted when the cell transforms into an amoeboid stage (Fig. 1R) or forms a cyst. Cysts are spherical and approximately 4–5 µm in diameter.

In the absence of uninfected algae, zoospores settle to the bottom, transform into amoebae, and eventually die. In contrast, zoospores attached to healthy algal cells retract the flagellum, form a cyst, and penetrate the host. Thus, the cyst stage is essential for infection.

Flagellar retraction occurs in two distinct ways: (1) gradual shortening of the flagellum until complete retraction (Fig. 1C, E), and (2) initial curling of the distal tip followed by retraction (Fig. 1F–H). In both cases, the flagellar axoneme relocates to the cell periphery and subsequently disassembles within the cytoplasm (Fig. 2B).

**Fig. 2.**
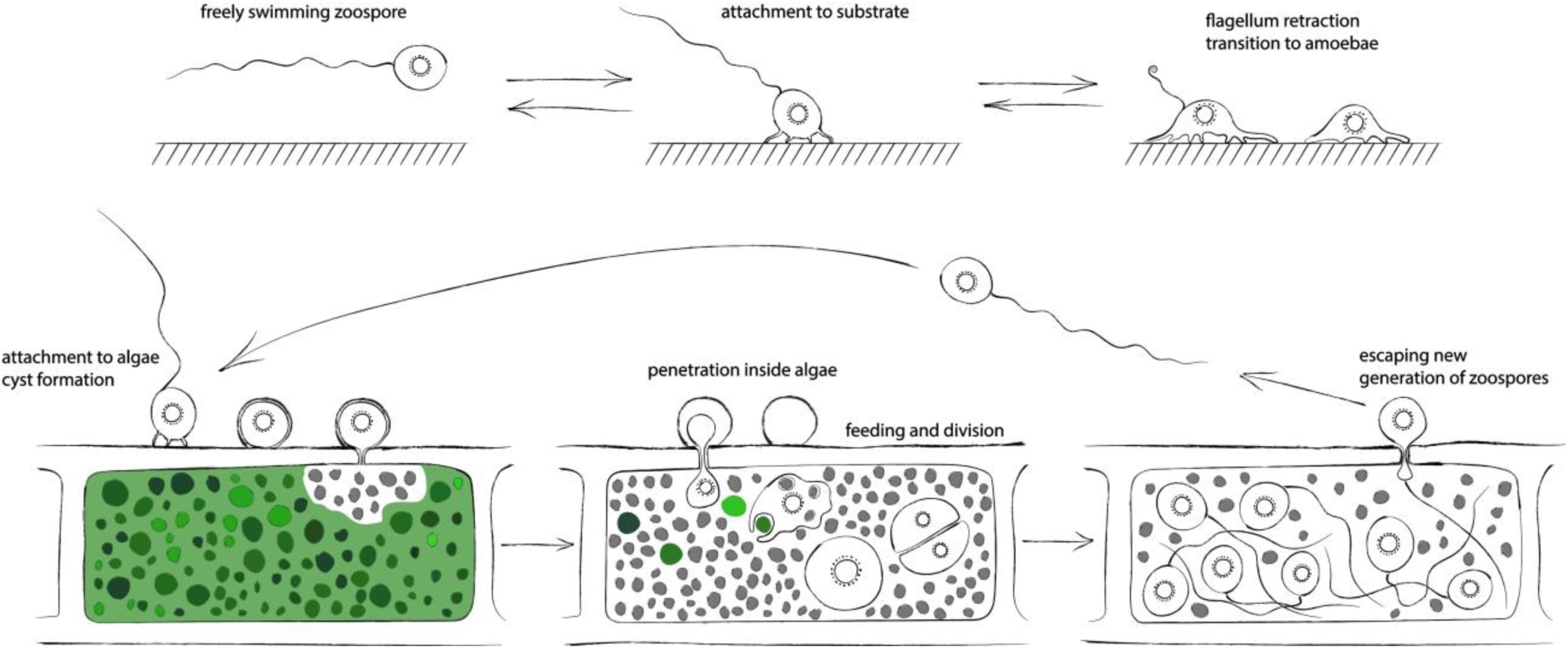
Drawing of *Insolitus nuclearius* life cycle. A – in the host absence. B – complete life cycle on the host *Dunaliella* sp.

Typically, numerous zoospores accumulate on the algal surface. During infection, two unusual and currently unexplained phenomena were observed (Video S1): (1) zoospores initially move erratically past a healthy green host cell without apparent recognition, but then suddenly converge toward it and attach, covering the cell surface (Fig. 1J); (2) shortly after encystment, the host cell contents become discolored, although phagocytosis has not yet begun and chloroplasts remain structurally intact. In general, infected host cells lose their green coloration and appear grey.

Following penetration, parasites phagocytize the algal contents, filling the entire space within the host cell wall. The cell interior contains spherical parasite cells and excretory material (Fig. 1L, M). The expulsion of large excretory bodies is illustrated in Fig. 1I. Each parasite cell appears to divide into two; this process was recorded in Video S2 and is shown in Fig. 1M.

Parasite cells can migrate into adjacent host cells by forming openings in the septa (Fig. 1N). For release, cells develop a flagellum and exit through the same opening in the algal wall used during penetration (Fig. 1O), or through a newly formed opening. SEM images of flagellates, amoeboflagellates, and amoebae bearing filopodia are shown in Fig. 1P–S.

A schematic representation of the life cycle is provided in Fig. 2.

### TEM observations

The spherical flagellated cells of strain X-141 are bounded by a plasma membrane (Fig. 3A, B). The vesicular nucleus is centrally located and surrounded by lipid globules and a single branched mitochondrion with lamellar cristae (Fig. 3A, B). At the posterior end of the cell, the mitochondrion is tightly adjacent to the nucleus and assumes a truncated conical shape, on the surface of which two kinetosomes are located (Fig. 3G; 4A). These kinetosomes are positioned on a common fibrillar plate adjacent to the mitochondrial surface (Fig. 3A, G–K). Dictyosomes of the Golgi apparatus are located on either side of the mitochondrial conical protrusion (Fig. 3A). A cluster of rough endoplasmic reticulum cisternae is often present at the anterior end of the cell (Fig. 3A). A highly flattened and branched microbody is also located adjacent to the anterior surface of the nucleus (Fig. 3B).

**Fig. 3.**
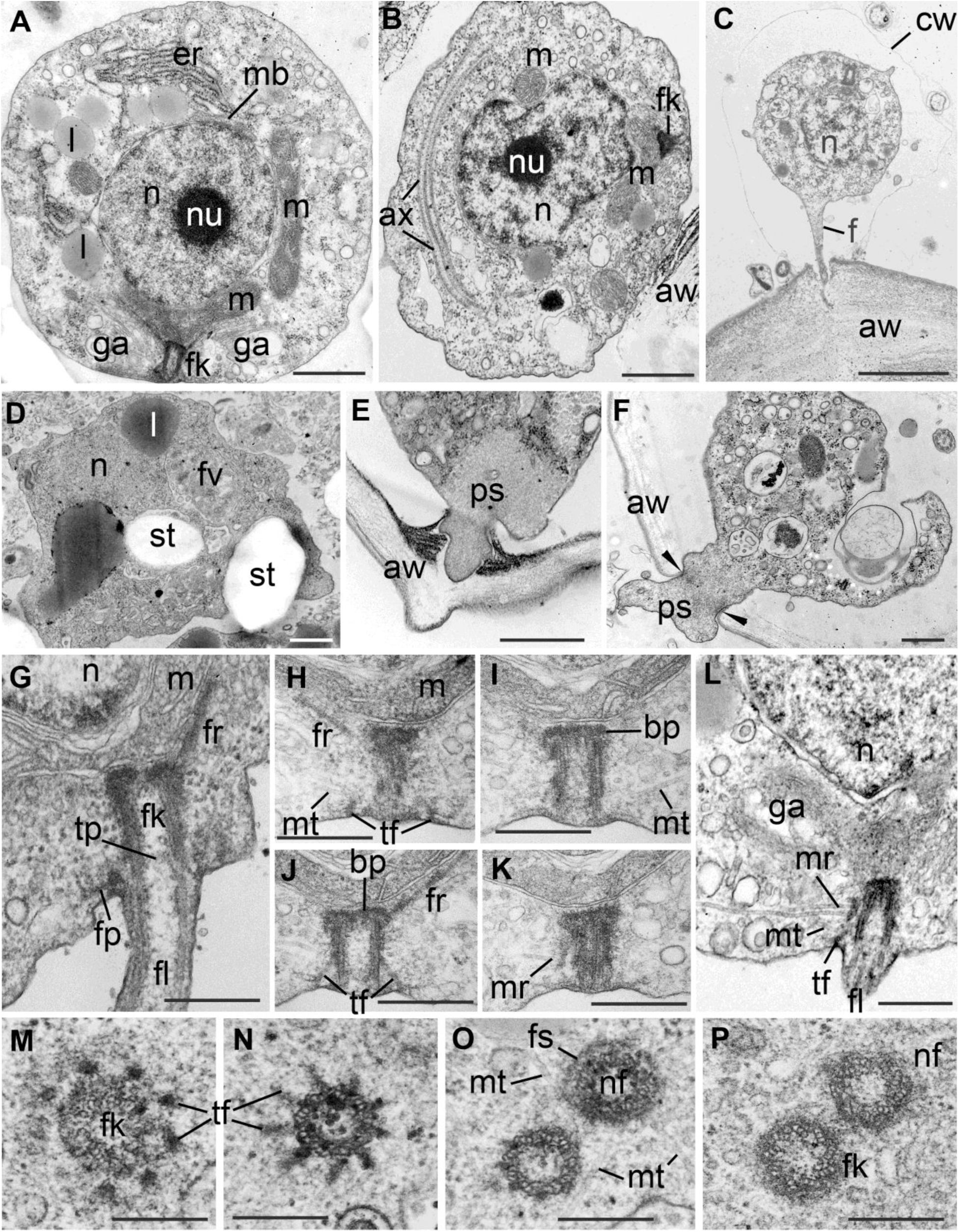
Ultrastructure of the main life cycle stages of *Insolitus nuclearius*. A – general view and organelle disposition in zoospore, C – two encysted zoospores on the algal surface, D – feeding cell inside the host with two starch granules excretes one of them (arrow), E – mature zoospore starts penetration the host cell wall using pseudopodium, F – zoospore release through a narrow pore in the host cell wall (arrowheads point desmosome-like contacts), G – serial longitudinal sections (LS) of flagellar kinetosome and transition zone, H-K – selected serial LS of kinetosome, L – slightly oblique section of kinetosome to show a lateral mt-root, M-P – selected serial transversal sections of flagellar and non-flagellar kinetosomes. Scale bars: A-B – 800 nm, C – 1 µm, D-P – 400 nm.

**Fig. 4.**
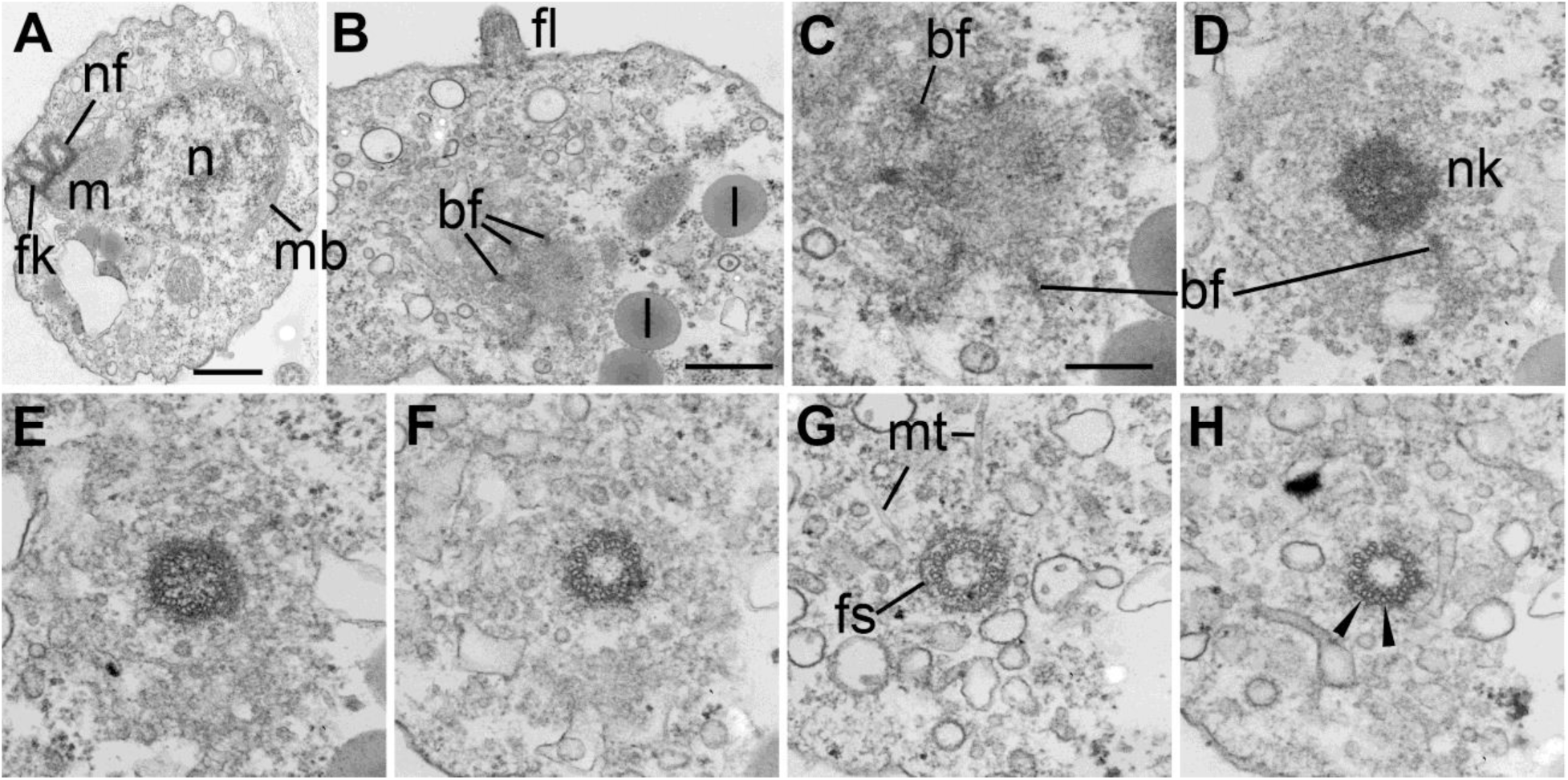
Flagellar apparatus structure of *Insolitus nuclearius*: A – LS of both kinetosomes, B shows a base of non-flagellar kinetosome apart from the base of flagellum. B-H – series of cross consecutive sections of non-flagellar kinetosome from its base (B,C) to the top (H). C is enlarged fragment of B. E shows a cartwheel structure at the proximal end, H shows microtubular doublets (arrowheads) at the distal end of kinetosome. Scale bars: A,B – 400 nm, C-H – 200 nm.

The kinetosomes are arranged either parallel to each other or at an acute angle (Figs. 3O, P; 4A). In rare sections, the kinetosomes appear separated, which may correspond to a specific stage of the cell cycle (Fig. 4B).

The non-flagellar kinetosome varies in length but is generally shorter than the flagellar kinetosome. Its distal region is surrounded by a ring of fibrillar material from which individual microtubules extend (Fig. 3O; 4G). The base of the non-flagellar kinetosome is attached to the mitochondrial surface by nine short fibrils, one originating from each microtubular triplet (Fig. 4B–D).

The flagellar kinetosome bears two cross-striated fibrillar roots at its proximal end, which extend along the mitochondrial surface (Fig. 3H, J). A fibrillar plaque associated with a lateral microtubular root is located closer to the distal end of the kinetosome (Fig. 3K, L). The flagellar kinetosome is characterized by prominent transitional fibers that extend approximately from the midpoint of each triplet and connect to the plasma membrane (Fig. 3J, M, N; 4A). From approximately the midpoint of each transitional fiber, one to three microtubules extend into the cytoplasm (Fig. 3H–K, M–P). The distal ends of the transitional fibers are attached to the plasma membrane via a distinct fibrillar lamina, which also gives rise to individual microtubules (Fig. 3G).

In the transition zone of the flagellum, only a broad terminal plate composed of low-density material is present at the distal end of the kinetosome (Fig. 3G). Upon flagellar loss, the transitional fibers are retained, and the entire kinetid detaches from the plasma membrane and moves into the cytoplasm. The complete kinetid–dictyosome–mitochondrion–nucleus complex is preserved in both the amoeboid stage and the cyst (Fig. 3B, C).

The cyst wall is thin and does not strictly maintain a spherical or oval shape (Fig. 3C). At the site of host contact, the cyst wall is absent, allowing the parasite plasma membrane to directly contact the algal cell wall and enabling the formation of a penetrating pseudopodium (Fig. 3C). The pseudopodium likely penetrates the algal wall mechanically by separating cellulose fibers to form a narrow channel; however, enzymatic activity (e.g., cellulases) may also be involved.

The intracellular feeding stages are characterized by dense cytoplasm, large lipid droplets, and digestive vacuoles containing plastid material and starch grains (Fig. 3D). After digestion of the host cytoplasmic contents, starch grains and other undigested residues are expelled from the parasite cell by exocytosis (Fig. 1O; 3D, F).

To exit the host cell or migrate to an adjacent cell through a septum, the amoeboid cell develops a flagellum and forms a specialized pseudopodium at its anterior end. The hyaloplasm of this pseudopodium consists of cytosol containing a dense network of actin filaments (Fig. 3E). This pointed extension likely separates the cell wall material and creates a passage sufficient for the entire parasite cell to pass through (Fig. 3F). At the contact site with the algal cell wall, desmosome-like junctions with electron-dense submembranous structures are formed, giving rise to bundles of microfilaments extending into the cytoplasm (Fig. 3F).

The ultrastructure of flagellated cells inside and outside the host does not differ significantly.

### Molecular phylogeny

The multigene phylogenetic tree was constructed based on 269 single-copy orthologs suitable for phylogenetic analysis of the Opisthokonta supergroup (Fig. 5). The tree was rooted using representatives of Amoebozoa as an outgroup. The overall branching topology of Holozoa and Holomycota is consistent with previously published phylogenies of Opisthokonta from recent studies (Tikhonenkov et al., 2020a; Galindo et al., 2022; Ocaña-Pallarès et al., 2022). Support values for all major clades are high, ranging from 90% to 100%.

**Fig. 5.**
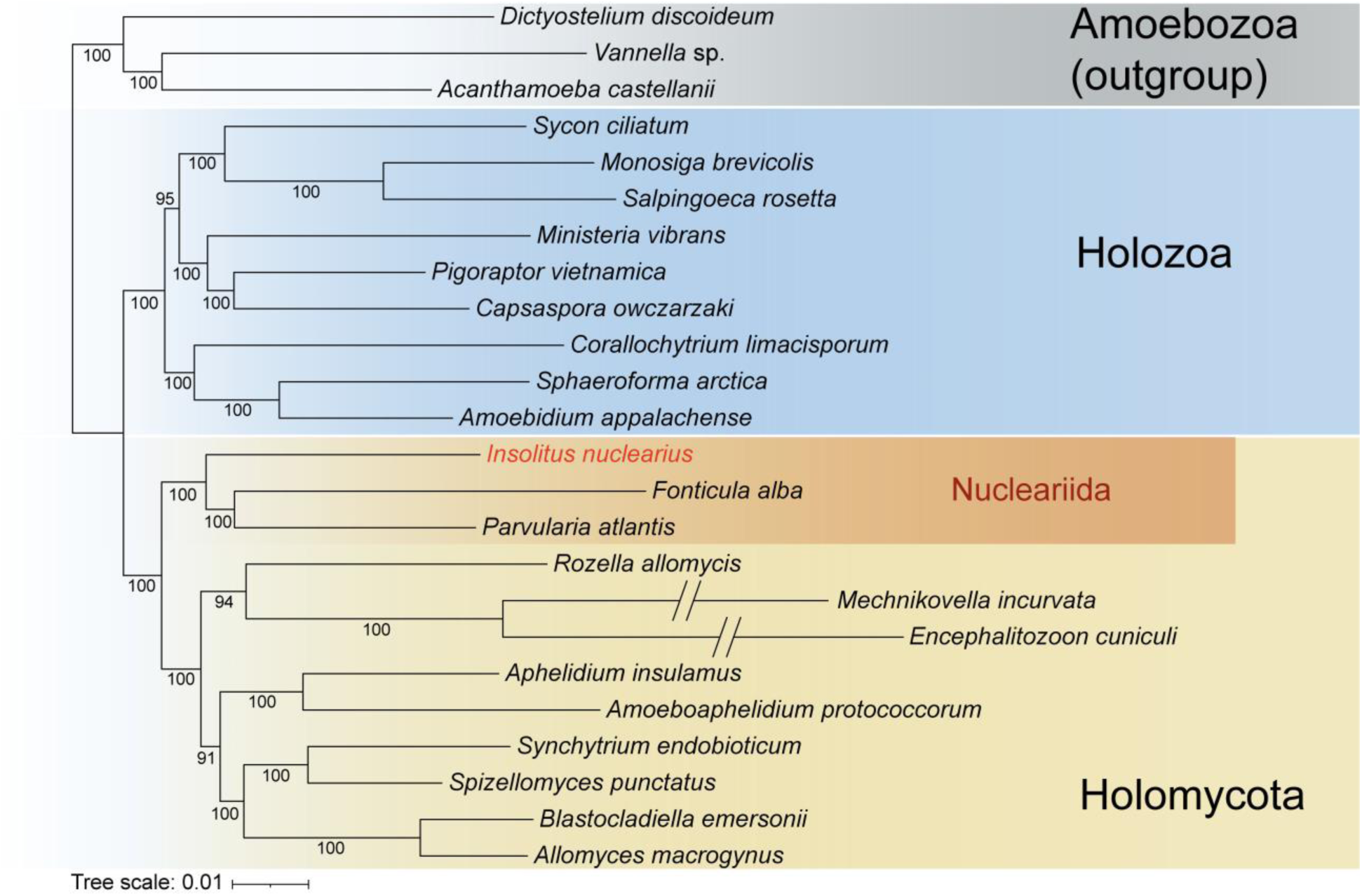
Phylogenetic tree of the Opisthokonta supergroup built based on 269 single-copy orthologs of 21 opisthokont and 3 amoebozoan species. The tree is rooted using amoebozoans as an outgroup. Phylogenetic branches of Amoebozoa, Holozoa, Holomycota and Nucleariida are highlighted in different colors. *Insolitus nuclearius* marked in red.

Strain X-141, described here as *Insolitus nuclearius* gen. et sp. nov. (see Taxonomic section below), belongs to Holomycota and is placed within the Nucleariida clade. This clade forms a sister group to all other Holomycota, including Rozellomycota (with Microsporidia), Aphelidiomycota, and Fungi, with full support. The studied organism appears as a sister lineage to *Fonticula* and *Parvularia*.

The ribosomal phylogeny supports these findings by placing the *Insolitus* lineage within nucleariids (Fig. 6). In contrast to the multigene analysis, the ribosomal tree positions *Insolitus* as a sister lineage to the clade comprising *Parvularia* and environmental OTUs, with moderate support. Together, *Insolitus* and *Parvularia* form a sister group to the *Nuclearia* clade.

**Fig. 6.**
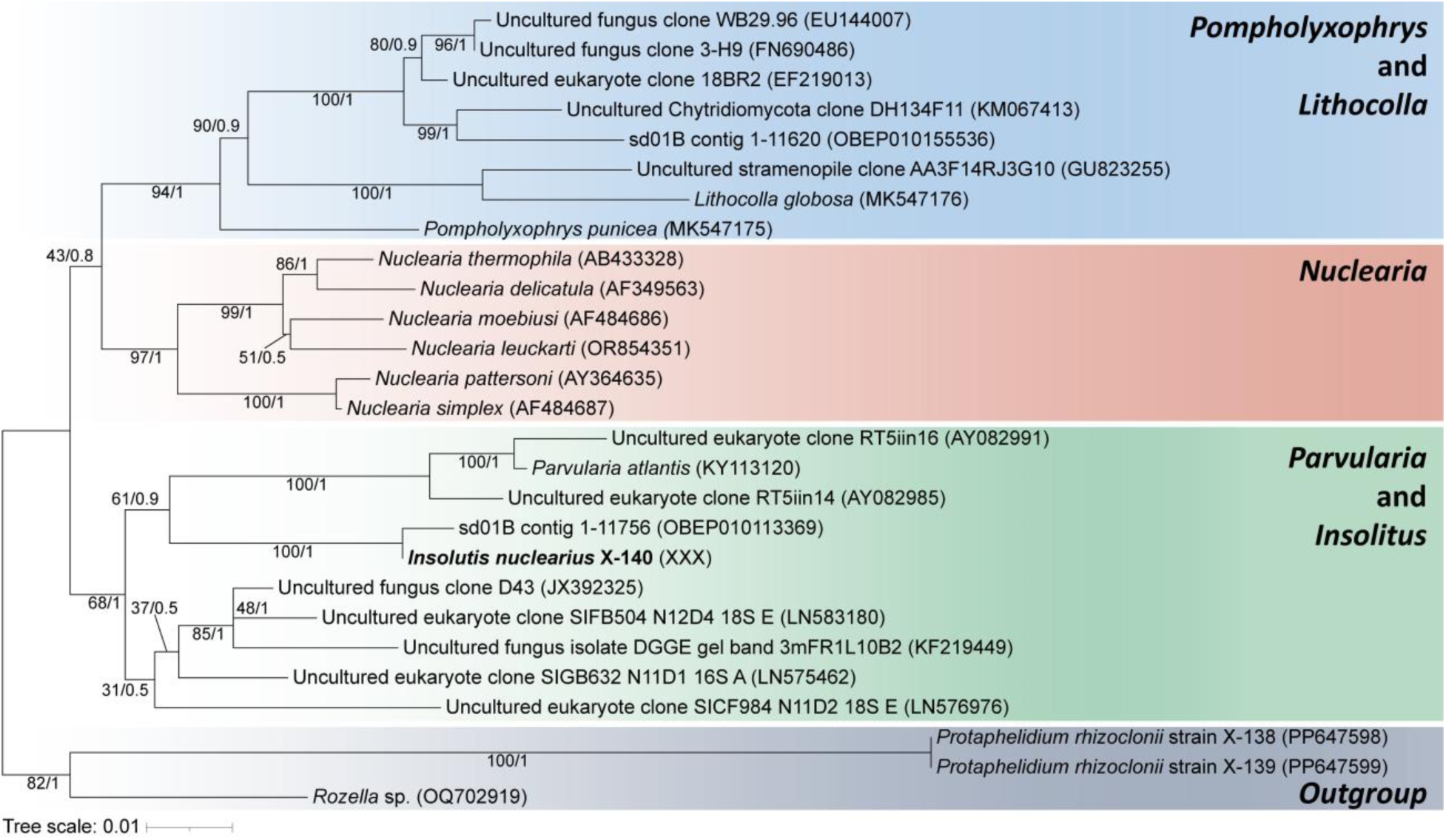
Maximum likelihood consensus tree of nucleariids and *Protaphelidium* with *Rozella* as outgroup based on 18S rDNA sequences, showing position of *Insolitus nuclearius* X-141 (red bold italic). The tree was constructed using 1363 nucleotide characters. Node support values are as follows: bootstrap values (IqTree) followed by Bayesian posterior probabilities (MrBayes).

The ribosomal sequence of the new representative is nearly identical to the environmental sequence OBEP010113369, which was previously detected in «sandy/muddy and dark sediment sample and collected in an area mostly covered by sea-salad, with water level 20-30 cm (water salinity is close to oceanic) and sampling depth 4-5 cm» in Limfjorden, Denmark (Karst et al., 2018).

## Discussion

### Phylogeny and taxonomy of strain X-141

When comparing the newly discovered representative of Nuclearida with other members of this group, only one clear similarity can be identified: the amoeboid trophic stage bearing filopodia resembles that of nucleariids, which are also capable of crawling within algal filaments and feeding on their contents, as observed in *Nuclearia thermophila* (Gabaldon et al., 2022). All other features of *Insolitus nuclearius* differ substantially, particularly its parasitic lifestyle and complex life cycle that includes a flagellated stage. Nevertheless, both multigene and ribosomal phylogenetic analyses robustly support its placement within Nuclearida. According to the multigene phylogeny, this strain represents a basal lineage within the group.

The process by which *Nuclearia* penetrates algal cells has not been described in detail, but it is likely that filopodia are used to create an opening in the cell wall and facilitate entry. We suggest that the amoeboflagellate stage of *I. nuclearius* may similarly use anterior filopodia to penetrate the algal cell wall during exit from or entry into host cells. We also do not exclude the involvement of GH5 endocellulases in this process, as demonstrated in the cercozoan *Orciraptor agilis*, which penetrates algal cells in a comparable manner (Moye et al., 2022).

Both Nuclearia species and viridiraptorids, when ingesting algal cytoplasm together with plastids, temporarily acquire coloration, which disappears following plastid digestion (Hess and Melkonian, 2013; Gabaldon et al., 2022). In contrast, *Insolitus nuclearius* appears to induce bleaching of chloroplasts even before penetration into the host cell, a phenomenon that requires further investigation. Furthermore, it does not appear to digest starch grains, which accumulate in the cytoplasm after plastid digestion and are subsequently excreted. This likely explains why both zoospores and starch grains are observed only within consumed algal cells.

The structure of the flagellated cell (zoospore) provides important information for comparison with other members of Holomycota and Holozoa. Several distinctive features were identified, including some that appear to be unique.

The curling of the distal end of the flagellum during retraction into the cell, observed during transformation into the amoeboid stage, has not previously been described.

The flagellar kinetosome is characterized by two basal fibrillar roots extending along the mitochondrial surface and a lateral microtubular root. In addition, microtubules originate from the midpoints of unusually large transitional fibers—an arrangement not reported in other protists. Similar large transitional fibers (referred to as “props”) have been described in zoosporic fungi (Barr, 1980, 2001), but the presence of microtubules arising from them, as well as their associated distal plate initiating microtubules, has not been documented. The base of the non-flagellar kinetosome is attached to the mitochondrial surface by nine fibrils, one from each triplet, which is also unprecedented among eukaryotes.

Such unusual features of the flagellar apparatus may reflect an ancestral state within Holomycota. Prominent transitional fibers are retained in zoosporic fungi, and the connection of kinetosomes to mitochondria via cross-striated fibrils has been described in *Rozella*, aphelids, and *Olpidium* (Karpov et al., 2019), indicating similarities between the kinetid of *I. nuclearius* and those of other flagellated holomycotans. However, the arrangement of two kinetosomes (flagellar and non-flagellar) positioned on the surface of a single mitochondrion with lamellar cristae is extremely rare. A comparable configuration is known in monoxenous trypanosomatids (Frolov, 2022), members of Excavata, which are considered among the most ancient eukaryotic lineages (Jewari and Baldauf, 2023).

Microbodies associated with the nucleus, often referred to as paranuclear bodies, are found in many groups of heterotrophic flagellates and amoebae and are typical of Chytridiomycota and Blastocladiomycota (Powell, 1978). Their functions are rarely discussed in morphological studies of protists; however, in fungi and aphelids they are involved in lipid metabolism and are typically located on the surface of lipid droplets or between lipid droplets and mitochondria (Powell, 2017a, b; Karpov et al., 2020).

According to ribosomal phylogeny, *I. nuclearius* belongs to a cluster that includes environmental sequences closely related to *Parvularia*. Species of *Parvularia* and other nucleariids generally feed primarily on bacteria, whereas Nuclearia species consume detritus, bacteria, and algae. However, *Nuclearia thermophila* has repeatedly been observed inside filamentous algal cells containing chloroplasts (Gabaldon et al., 2022). Even the original description of *N. delicatula* reported its tendency to penetrate algal cells using pseudopodia to obtain nutrients (Cienkowski, 1865). Thus, nucleariids possess the ability to penetrate algal cells and feed on their contents, which may represent a preadaptation to intracellular parasitism.

This type of feeding strategy (predation) is common among Cercozoa (e.g., *Orciraptor agilis*), where a transition from predation to parasitism is also observed, as in *Viridiraptor invadens* (Hess and Melkonian, 2013), as well as in the amoebozoan *Idionectes* (Hess, 2021), in which the organism not only consumes algal contents but also reproduces within the host cell.

The life cycle of *Insolitus nuclearius* is superficially similar to that of parasitic protists; however, its phylogenetic position and its morphological and biological characteristics are highly distinctive, justifying its designation as a new genus and species within Nuclearida.

Our discovery of this highly unusual representative of Nuclearida necessitates a revision of the taxonomy of the group.

First, Nuclearida are not exclusively free-living amoebae but also include flagellated parasitic forms. Therefore, the diagnoses of the order Nucleariida, class Nuclearidea, and phylum Nuclearida should be emended.

Second, ribosomal phylogenetic analyses reveal three main clusters (Galindo et al., 2019; Gabaldon et al., 2022; present study, excluding *Fonticula*), which correspond to distinct morphological groups: (1) the clade including flagellated *Insolitus nuclearius* together with *Parvularia* and *Fonticula* represents the most basal lineage; (2) *Nuclearia* includes amoebae covered with a mucous coat; and (3) *Lithocolla* and *Pompholyxophrys* comprise organisms with a rigid covering composed of inorganic particles. These groups could potentially correspond to separate families. However, this division is not consistently supported across ribosomal phylogenies, where many nodes show low support and the position of *Fonticula* remains unstable. In contrast, multigene phylogeny supports division into two families, and we therefore provisionally recognize two families within Nucleariida.

Third, Nuclearida represents a monophyletic lineage within Holomycota and should be assigned a taxonomic rank comparable to that of Rozellomycota and Aphelida, i.e., at least at the phylum level. Tedersoo et al. (2018) proposed the kingdom Nucleariae, comprising two phyla, Nuclearida and Fonticulida, with corresponding classes Nuclearidea and Fonticulea. At present, however, the assignment of nucleariids to kingdom rank, as well as the status of Fonticula at the phylum and class levels, remains uncertain. Multigene phylogenies place *Fonticula* together with *Parvularia*, while ribosomal trees alternatively associate it with *Nuclearia* or *Parvularia* (Galindo et al., 2019; Gabaldon et al., 2022). This suggests that *Fonticula* should be included within the family Nucleariidae.

### Taxonomy of Nuclearida

#### Phylum Nuclearida (Tedersoo et al.), emend. Karpov, Pozdnyakov & Seliuk

Index Fungorum ID:

Diagnosis: Free-living or parasitic, uni- or multinucleate filose amoebae; cells typically covered by a mucous coat; life cycle of parasitic representatives includes an intracellular, feeding filopodial amoeba and an amoeboflagellated dispersal stage; mitochondria with flat or discoid cristae, from two to multiple dictyosomes, a microbody, lipid inclusions, and cytoskeletal elements composed of microtubules and fibrils associated with flagellar apparatus. The flagellar apparatus consists of flagellar and non-flagellar kinetosomes connected to mitochondrial surface; flagellar kinetosome bears a lateral microtubular root and prominent transitional fibers producing microtubules; cysts unknown. In multigene phylogenies, Nuclearida form a well-supported clade sister to all other Holomycota.

Type genus: *Nuclearia* Cienkowski, 1865.

Class Nuclearidea (Tedersoo et al.), emend. Karpov, Pozdnyakov & Seliuk

Index Fungorum number:

Diagnosis: As for the phylum.

#### Order Nucleariida (Cavalier-Smith), emend. Karpov, Pozdnyakov & Seliuk

Index Fungorum number:

Diagnosis: Free-living or parasitic filose amoebae, spherical to flattened, with a vesicular nucleus; uni- or multinucleate. Free-living species exhibit two stages: a vegetative filopodial amoeba covered by a mucous coat and a cyst. Parasitic representatives possess a life cycle including an intracellular feeding amoeba, an amoeboflagellated dispersal stage, and an infective cyst. Radiating filopodia, sometimes branching, tapering when attached to substrates, and lack extrusomes. Mitochondria have flat or discoid cristae. Cells contain two to multiple dictyosomes, microbodies, lipid inclusions, and cytoskeletal elements associated with the flagellar apparatus. The flagellar apparatus consists of flagellar and non-flagellar kinetosomes located on the mitochondrial surface and connected to it by fibrillar roots. The flagellar kinetosome bears a lateral microtubular root and well-developed transitional fibers producing microtubules. Cell size ranges from nanoplanktonic to small microplanktonic forms. Members form a robust clade sister to other holomycotans.

Note: As a formal diagnosis of the order was not provided in Cavalier-Smith (1993), the present diagnosis is based on the comprehensive description by Gabaldon et al. (2022).

#### Family Insolitidae Karpov, Pozdnyakov & Seliuk, fam. nov

Index Fungorum number:

Diagnosis: Nucleariids possessing an amoeboflagellate dispersal stage (zoospore). The zoospore transforms into an amoeboflagellate that moves using branching filopodia and a non-motile posterior flagellum. The flagellum is retracted as the cell transforms into an amoeba and subsequently forms a cyst. The cyst penetrates the algal cell wall, enters the host, and phagocytizes the cellular contents. The parasite divides into two daughter cells, which develop a posterior flagellum and exits through an opening in the algal wall. The structure of the flagellar apparatus corresponds to that of the order.

#### Genus *Insolitus* Seliuk & Karpov, gen. nov

Index Fungorum number:

Etymology: From Latin *insolitus*, meaning “unusual”.

Type species: *Insolitus nuclearius* Seliuk & Karpov, sp. nov.

Diagnosis: Amoeboflagellate zoospores bearing a single posterior opisthokont flagellum and anterior filopodia. The zoospore attaches to a host cell, encysts, penetrates the host as an amoeba, phagocytizes host contents, and divides into two daughter cells. Each daughter cell develops a flagellum and exits the host through an opening in the cell wall. The amoeboflagellate possesses two nearly parallel kinetosomes located on the surface of a mitochondrion with flat cristae, positioned posterior to the nucleus; a flattened microbody is located anterior to the nucleus. The flagellar kinetosome bears two cross-striated fibrillar roots, a lateral microtubular root, and prominent transitional fibers that give rise to microtubules. The non-flagellar kinetosome has nine fibrillar basal elements and a fibrillar sheath producing individual microtubules. Parasites of marine green algae.

#### *Insolitus nuclearius* Seliuk & Karpov, sp. nov.: Fig. 1-3, present paper

Index Fungorum number:

Etymology: From Latin *nuclearius*, referring to its affinity with *Nuclearia*.

Diagnosis: Spherical zoospores 4–5.5 μm in diameter, bearing a posterior flagellum up to 22 μm in length. Numerous filopodia are present and may be long and branching. Spherical cysts (4–5 μm in diameter) penetrate the host cell wall and phagocytize host contents.

Habitat: Marine; isolated from coastal green algae.

Host: Marine green alga *Ulva*.

Type locality: Russia, White Sea, near Poyakonda village (66.5549° N, 33.09907° E), collected in August 2023 by A.O. Seliuk. Type material deposited as epoxy resin blocks (accession numbers X-141-Pr6–Pr8) in CCPP ZIN RAS. GenBank accession: SSU XXX.

Additional material: Isolate K-1 from a sample collected at the same time approximately 700 m from the type locality; isolates Kar-9, Kar-10, and Kar-11 from samples collected in June 2025 near the ZIN RAS biological station “Kartesh”. These isolates have not yet been confirmed by molecular phylogenetic analysis.

Note: *Insolitus nuclearius* differs fundamentally from all previously described nucleariids, both naked and scale-bearing forms. It also exhibits elements of coordinated (social-like) feeding behavior.

Family Nucleariidae Cann and Page, 1979

Uni- or plurinucleate, somewhat flattened or spherical amoebae covered with mucilage, siliceous scales (idiosomes), or xenosomes; moving by attachment and subsequent shortening of filopodia.

Confirmed genera: *Nuclearia* Cienkowsky, 1865, *Parvularia* López-Escardó et al. 2018, *Fonticula* Worley et al. 1979, *Pompholyxophrys* Archer, 1869 and *Lithocolla* Schulze, 1874.

### Ancestor of Holomycota

In multigene phylogenies that do not include additional nucleariid representatives, *Insolitus nuclearius* appears as a sister lineage to *Parvularia* and *Fonticula*. Broader phylogenomic analyses likewise support the grouping of *Parvularia* and *Fonticula* (Galindo et al., 2019). Considering that these lineages are followed by *Nuclearia* and, subsequently, by the scale-bearing *Lithocolla* and *Pompholyxophrys* in ribosomal phylogenies (Fig. 5; Gabaldon et al., 2022), *Insolitus* occupies the most basal position among nucleariids. This suggests that it may retain characteristics close to those of the ancestral nucleariid.

This observation necessitates a revision of the evolutionary scenario proposed by Gabaldon et al. (2022), which assumes that *Parvularia* and *Fonticula* represent the most basal lineage of Holomycota. Feeding on algae requires either engulfment of small cells or penetration of larger algal cells to access their cytoplasm. Nucleariid amoebae are capable of both strategies: engulfing bacteria and small algae or penetrating larger algal cells.

We propose that the last common ancestor (LCA) of nucleariids was a small, naked marine flagellate feeding on bacteria and small algae. This organism was able to form filopodia and retract its flagellum to transition into an amoeboid stage, which could also encyst. Thus, it likely exhibited alternating motility using both the flagellum and filopodia and flexibility in feeding strategies.

This ancestral form may have diversified into the known nucleariid lineages. Through its capacity for morphological transformation, it could have adapted to feeding on larger algae with filamentous or thalloid structures, as observed in *Insolitus*. The emergence of a free-living amoeboid stage may have facilitated the development of a mucous coat, as seen in *Parvularia* and *Nuclearia* (Galindo et al., 2019), which could subsequently have enabled the incorporation of exogenous materials (as in *Lithocolla*) or the secretion of endogenous siliceous elements (as in *Pompholyxophrys*) (Gabaldon et al., 2022).

Nuclearia may have independently acquired the ability to exploit algae as a food source, and in filamentous algae this could have led to internal feeding via penetration using filopodia (Gabaldon et al., 2022). *Fonticula* differs from other nucleariids in exhibiting aggregative multicellular behavior during the formation of fruiting bodies (Toret et al., 2022). A comparable, although less pronounced, aggregation behavior was observed in *Insolitus* during feeding.

Lineages of Holomycota such as Rozellida and Aphelida exhibit nutritional strategies similar to that of *Insolitus*. Their trophic stages are endobiotic but remain phagotrophic. In the ancestor of Fungi, such trophic forms may have later transitioned to saprotrophic feeding on the cell walls of dead algae (Chang et al., 2015). The amoeboflagellate stage may have been reduced to a dispersal non-feeding zoospore, while the filose amoeboid stage was lost.

### Ancestor of Holozoa and Opisthokonta

The ancestor of basal Holomycota lineages is also the common ancestor of Holozoa (Fig. 7). Recent studies suggest that the earliest-branching holozoan lineage is Ichthyophonida, which includes amoeboid parasites of invertebrates (Tikhonenkov et al., 2020a, b; Aleoshin et al., 2026). Among them, only *Dermocystidium percae* possesses a dispersal stage with a single flagellum, accompanied by a non-flagellar orthogonal kinetosome and a cross-striated fibrillar root (Pekkarinen et al., 2003). This structure is much simpler than the kinetid observed in *Insolitus* and lacks a connection to the mitochondrion.

**Fig. 7.**
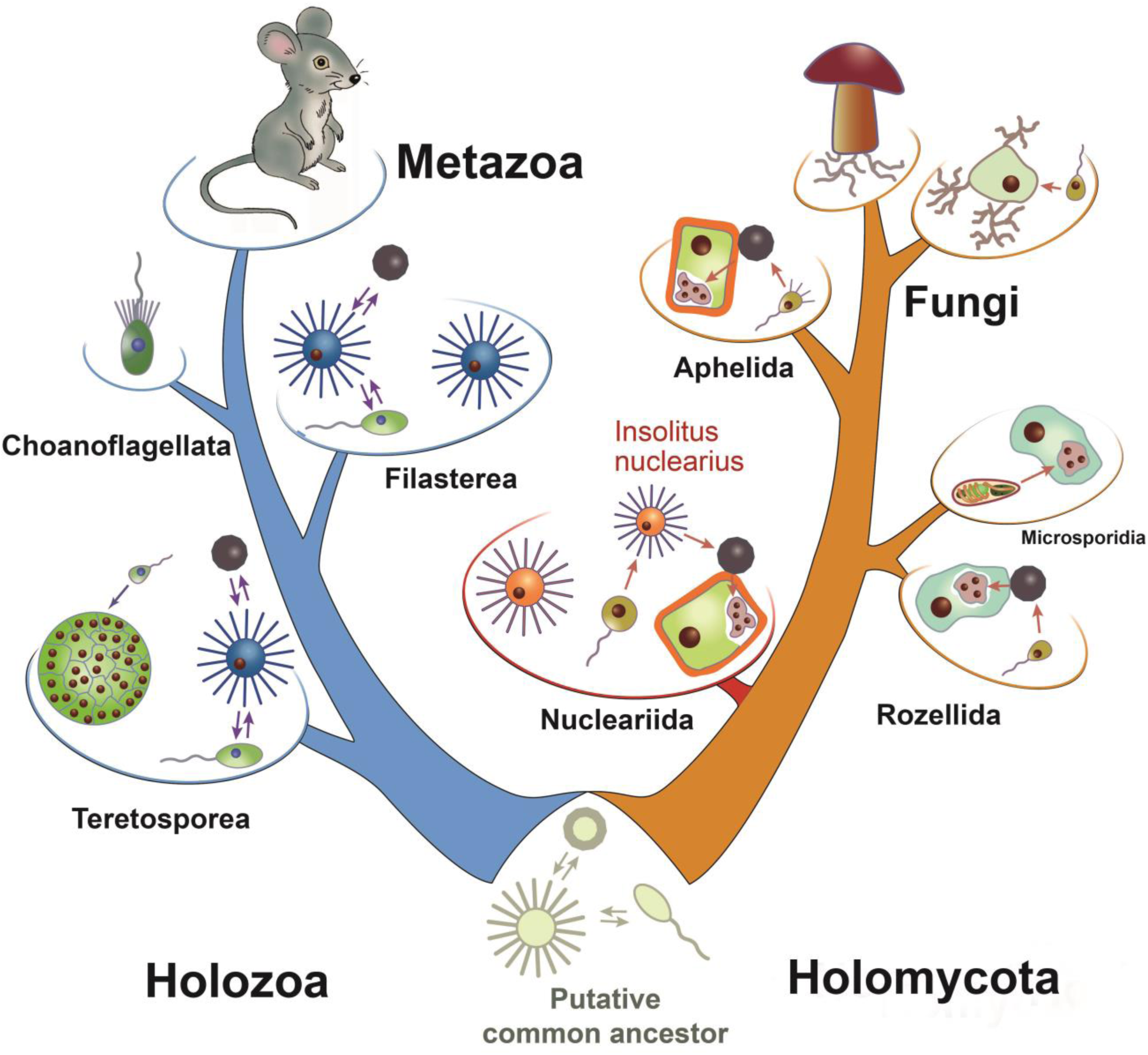
Schematic drawing of opisthokont phylogeny of Holozoa and Holomycota evolved from putative ancestor. Key stages of life cycles of opisthokont representatives and its ancestor are illustrated. *Insolitus nuclearius* is designated by the genus name.

A more complex kinetid is found in the next holozoan lineage, Pluriformea, which includes the uniflagellated predator *Syssomonas* (Hehenberger et al., 2017; Tikhonenkov et al., 2020a). *Syssomonas multiformis* exhibits flagellated, amoeboid, and cyst stages and possesses flat mitochondrial cristae; however, its kinetid differs markedly from that of *Insolitus*. The flagellar and non-flagellar kinetosomes are neither parallel nor connected to the mitochondrion or nucleus. The transition zone is longer and includes both terminal and transverse plates. Transitional fibers are of standard size and do not give rise to microtubules, and the non-flagellar kinetosome lacks a fibrillar sheath. The only comparable feature is the presence of dense material at the kinetosomal surface producing lateral microtubules, although this structure is more prominent in *Syssomonas* and forms a radial array reminiscent of choanoflagellates (Karpov and Leadbeater, 1998).

Within Filasterea, primarily composed of free-living filopodial amoebae, two flagellated forms are known: *Pigoraptor* and *Ministeria* (Tikhonenkov et al., 2020a; Torruella et al., 2015; Mylnikov et al., 2019). Both possess orthogonal kinetosomes located near the nucleus, and in *Pigoraptor* these give rise to disorganized microtubules extending into the cytoplasm. It has been suggested that elongation of the transition zone in *Syssomonas*-like ancestors may have led to the development of a central filament, as observed in *Pigoraptor*. This type of transition zone, first described in choanoflagellates (Hibberd, 1975; Karpov, 1982, 1985, 2000), is now characteristic of both Pluriformea and choanoflagellates (Tikhonenkov et al., 2020a).

Another flagellated holozoan, *Tunicaraptor*, occupies an uncertain phylogenetic position, either as a sister group to filasterians or as the earliest-branching holozoan (Tikhonenkov et al., 2020b). *Tunicaraptor unikontum* is a marine predator with an anterior feeding structure (“mouth”) and a posterior flagellum. Its flagellar apparatus includes flagellar and non-flagellar kinetosomes arranged orthogonally or at an acute angle. The flagellar kinetosome bears a lateral microtubular root, similar to that of *Syssomonas*. Overall, holozoan flagellates exhibit relatively simple root systems, evolving from lateral microtubular roots to the radial microtubule arrays characteristic of choanoflagellates.

Thus, the flagellar apparatus of holozoans differs fundamentally from that of *Insolitus*, supporting its placement within Holomycota. At the same time, *Insolitus* shares with holozoans the presence of a fibrillar structure associated with the kinetosome that gives rise to lateral microtubules. This feature may have been present in the kinetid of the opisthokont last common ancestor.

The life cycles of *Insolitus* and unicellular holozoans share a high degree of polymorphism. In holozoans, parasitism likely evolved secondarily, as their hosts are multicellular animals that appeared later in evolution. No holozoans are known to parasitize unicellular organisms, suggesting that their ancestor was free-living. Most unicellular holozoans exhibit polymorphism, transitioning between flagellated and amoeboid forms.

A similar polymorphic ability of *Insolitus* may reflect an ancestral state inherited from the opisthokont ancestor. This ancestor was likely a free-living, polymorphic protist capable of cytoplasmic feeding, with both flagellated and amoeboid stages and a resting cyst stage. Divergence between Holomycota and Holozoa may have been driven by differences in feeding strategies and preferred prey. One lineage retained predominantly motile flagellated predators, whereas the other developed the ability to penetrate rigid cell walls, with an amoeboid trophic stage and a flagellated dispersal stage.

Thus, the newly discovered nucleariid represents a key transitional form that integrates evolutionary histories of Holomycota and the broader Opisthokonta supergroup and provides new insights into their ancestral states and early diversification.

## Acknowledgements

The authors acknowledge RSF grant 26-14-00265 and Saint Petersburg State University for access to the equipment of the core facility center RMiCT of the Research Park SPbSU.

